# Further expansion of methane metabolism in the Archaea

**DOI:** 10.1101/312082

**Authors:** Yulin Wang, Zheng-Shuang Hua, Kian Mau Goh, Paul N. Evans, Lei Liu, Yanping Mao, Philip Hugenholtz, Gene W. Tyson, Wen-Jun Li, Tong Zhang

## Abstract

The recent discovery of key methane-metabolizing genes in the genomes from the archaeal phyla *Bathyarchaeota* and *Verstraetearchaeota* has expanded our understanding of the distribution of methane metabolism outside of the phylum *Euryarchaeota*. Here, we recovered two near-complete crenarchaeotal metagenome-assembled genomes (MAGs) from circumneutral hot springs that contain genes for methanogenesis, including the genes that encode for the key methyl-coenzyme M reductase (MCR) complex. These newly recovered archaea phylogenetically cluster with *Geoarchaeota* (deep lineage of archaeal order *Thermoproteales)*, and the MCR-encoding genes clustered with the recently reported methanogens within the *Verstraetearchaeota*. In addition, genes encoding hydroxybutyryl-CoA dehydratase were identified in the newly recovered methanogens, indicating they might carry out the β-oxidation process. Together, our findings further expanded the methane metabolism outside the phylum *Euryarchaeota*.

## Introduction

Methane-metabolizing archaea play a key role in the global carbon cycle as they are major contributors to the formation or oxidation of methane (Reeburgh 2007; Thauer *et al*., 2008). The recent discoveries of predicted methanogenic archaea in the phyla *Bathyarchaeota* (Evans *et al*., 2015) and *Verstraetearchaeota* (Vanwonterghem *et al*., 2016) challenged our original hypothesis that methane metabolism originated from the phylum *Euryarchaeota* (Gribaldo and Brochier-Armanet 2006), indicating the current phylogenetic distribution of methanogens or archaeal methanotrophs will be expanded. The methyl-CoM reductase (MCR) complex not only act as a key role in the methanogenesis, but also activate methane to methyl-coenzyme M in anaerobic methane oxidation process (Knittel and Boetius 2009; Orphan *et al*., 2001). However, a recently detected *Ca. Syntrophoarchaeum* organisms that was reported to catalyze propane and butane oxidation via enzymes similar to methyl-coenzyme M reductase (Laso-Perez *et al*., 2016), indicating the MCR complex might not be limited to the activation of methane in archaea. The similarity of *Ca. Syntrophoarchaeum* and *Bathyarchaeota* MCR sequences suggests the *Bathyarchaeota* might play a role in short-chain hydrocarbons oxidation. These results indicate the mechanisms associated methane metabolism in archaea requires further study. In this study, culture-independent metagenomics was used to recover previously unrecognized methane-metabolizing archaea present in hyperthermal hot springs.

## Recovery of crenarchaeotal genome with divergent MCR complex

In this study, we collected biomass from two different spring heads of Ulu Slim hot spring (UShs), Malaysia. The temperatures of two spring heads were 89 °C (US89) and 80 °C (US80). A total of 43 metagenome-assembled genomes (MAGs) were recovered from the co-assembled metagenome (US89, ~36 Gb; US80, ~36 Gb), including 1 *Crenarchaeota-like* (Cren_UShs, completeness of 93.4%) MAG, which contains genes encoding for the MCR complex *(mcrABG)* and accessory proteins (*mcrCD*). One additional *mcrABG* complex was detected from a contig (C_19902, 16.8 kb) that was not recruited to any MAG. Using these newly recovered *mcrA* gene sequences, we recovered several closely related *mcrA* sequences from two hot spring metagenomes: Jinze hot spring (JZhs), Tengchong, China (Eloe-Fadrosh *et al*., 2016) and Great boiling hot spring (GBhs), Nevada, United States of America (Peacock *et al*., 2013). Another near complete (completeness of 99.0%) *Crenarchaeota-like* MAG that contains genes that encode for a complete MCR complex in JZhs (Cren_JZhs, Supplementary Figure S1 and Table S1) was recovered.

These newly recovered *mcrA* genes have a close relationship with homologues in recently reported *Verstraetearchaeota* genomes (70-90% amino acid (aa) identity). Phylogenetic analyses of the McrA amino acid sequences confirmed the newly recovered *mcrA* gene sequences are basal to the *Verstraetearchaeota* homologues and form a cluster distant to the *Euryarchaeota, Bathyarchaeota* and methanotrophic McrA clades (Fig. 1a). The close relationship with *Verstraetearchaeota* was also confirmed by the phylogenetic analysis of McrB (Supplementary Figure S2) and McrG (Supplementary Figure S3). However, the low average genomic amino acid identity (AAI, 45.9-48.4%, Supplementary Table S2) and variable co-located gene loci (Fig. 1b) between the newly recovered MAGs and *Verstraetearchaeota* genomes suggest Cren_UShs and Cren_JZhs are distinct to *Verstraetearchaeota*. A genome tree (Fig. 2a) constructed using 56 concatenated ribosomal proteins (RPs) of 704 archaeal genomes place the Cren_UShs and Cren_JZhs genomes adjacent to the *Geoarchaeota* (Kozubal *et al*., 2013) which is a deep lineage of the archaeal order *Thermoproteales* (Guy *et al*., 2014). Furthermore, the tetra-nucleotide word frequency pattern confirms the Cren_UShs and Cren_JZhs are divergent from members in the phylum *Verstraetearchaeota* (Fig. 2b and 2c). These phylogenetic and genome attributes confirm that the newly recovered MAGs are previously unrecognized crenarchaeotal archaea that are likely associated with methane-metabolism.

**Figure 1.**
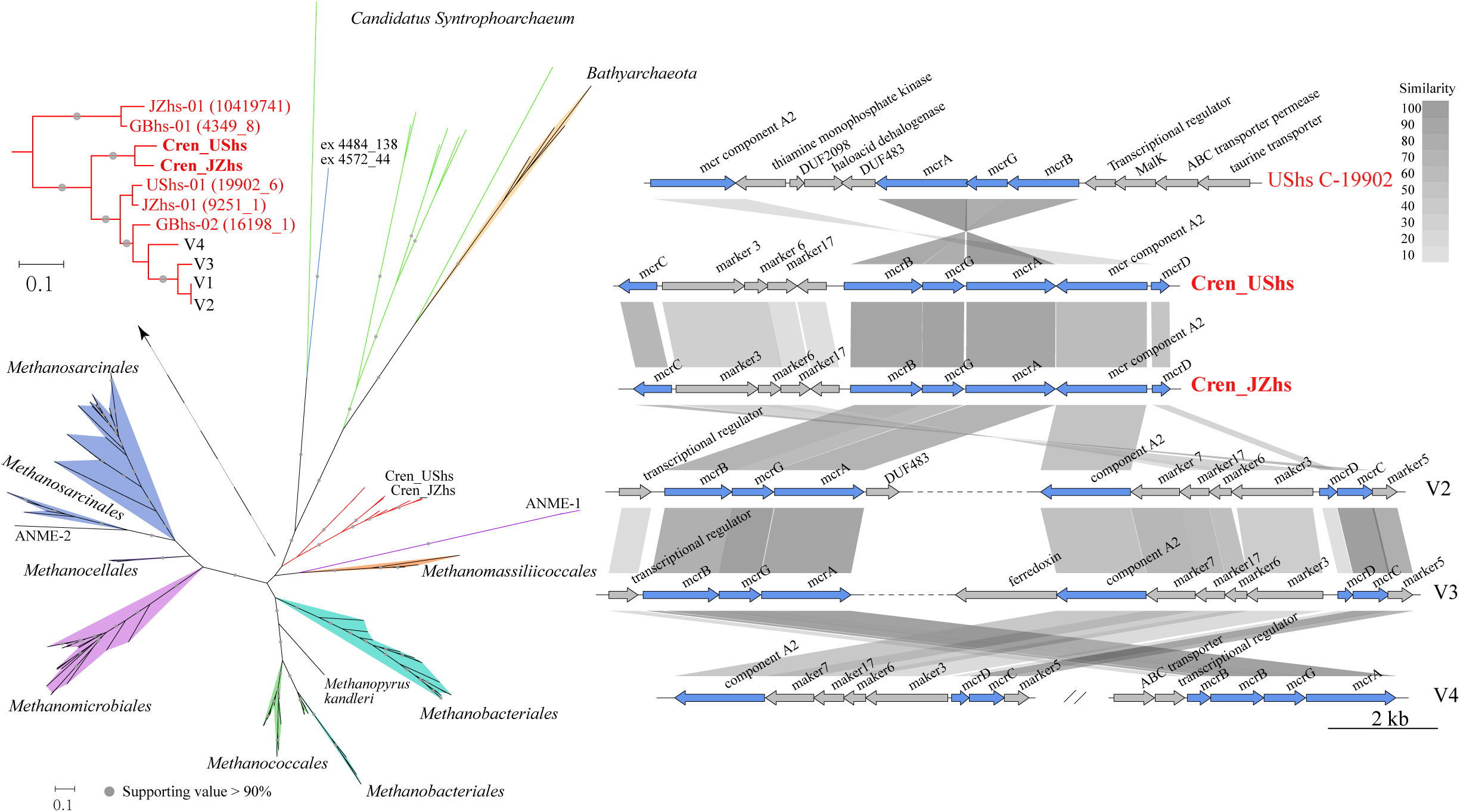
Phylogeny of mcrA gene and gene loci of the contigs/scaffolds that carry MCR complex. a, McrA amino acid sequences of newly recovered crenarchaeotal methanogens and reported methanogenic organisms were used to construct the phylogenetic tree. b, Homologous similarity was calculated using Blastp, the dotted black lines indicate continuity loci where unrelated genes are not displayed, and parallel double lines symbolize a break in locus organization (different contigs). MCR complex was colored in blue. Genes and noncoding regions are drawn to scale.

**Figure 2.**
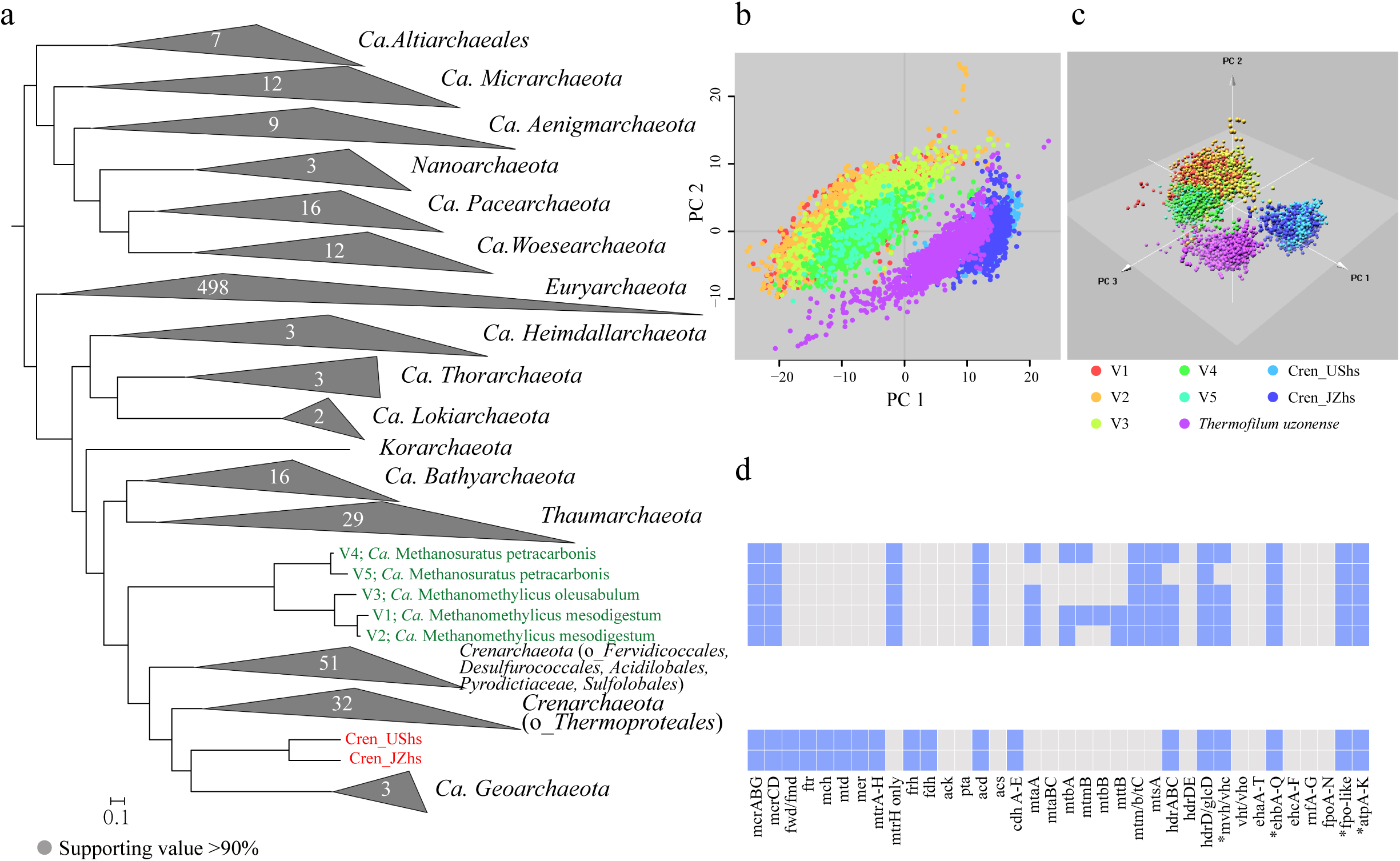
Genome comparison based on genome tree, sequence composition and distribution of genes involved in methane metabolism. a, Genome tree of 56 concatenated syntenic ribosomal proteins from 704 archaeal genomes, including two MAGs (Cren_UShs and Cren_JZhs) recovered in this study. b, Scaffolds/chromosome of selected genomes were shredded into sequences with length of 5 kb (4 kb overlapping). to perform the tetra-nucleotide word frequency PCA. c, Presence (blue)/absence (grey) of genes involved in methane metabolism and energy conservation for the newly recovered MAGs and reported Verstraetearchaeota (V1-V5) genomes. Annotation was based on the IMG/ER and KEGG GhostKOALA. Asterisks indicate not all subunits are identified in Cren_UShs and Cren_JZhs.

## Methane metabolism

KEGG orthology (KO) based analysis revealed the metabolic features of Cren_UShs and Cren_JZhs are highly similar to the *Verstraetearchaeota* and *Bathyarchaeota* genomes and distant to euryarchaeotal methanogens and methanotrophs (Supplementary Figure S4). Also, these newly recovered MAGs share more gene orthologues with *Verstraetearchaeota* populations (V1-V4, 566) than with *mcr-*containing *Bathyarchaeota* populations (BA1 and BA2, 225) (Supplementary Figure S5a), which was in line with the phylogenetic analyses of McrA. Only 160 and 156 gene orthologues were found to be unique in Cren_UShs and Cren_JZhs genomes, respectively, when compared to V2-V4 representatives of the phylum *Verstraetearchaeota* (Supplementary Figure S5b), indicating some different metabolic pathways have potentially been adopted by this newly recovered crenarchaeotal methanogens.

In addition to the key genes coding for MCR complex *(mcrABG)*, core methane metabolism pathway genes in Cren_UShs and Cren_JZhs (Fig. 2d) resemble the hyperthermophilic hydrogenotrophic methanogen *Methanopyrales kanderli* (Slesarev *et al*., 2002). However, *mcr* gene and genome trees place these organisms basal to *Verstraetearchaeota* and other H_2_-depdendent methylotrophs, suggesting these methylotrophs may have lost the ability for hydrogenotrophic methanogenesis (Borrel *et al*., 2016), while the hyperthermophilic *Crenarchaeota-like* organisms have retained this function (Fig. 3). Most of genes encoding methyl-H_4_MPT S-methyltransferase (Mtr) were not found in *Verstraetearchaeota* and *Bathyarchaeota*. In contrast, *mtrABCDEFGH* genes are all found in these two MAGs (Supplementary Table S3). The remaining genes for the hydrogenotrophic methanogenesis or anaerobic methane oxidation are identified in the present study, which are missing from the methane-metabolizing archaea within *Verstraetearchaeota* and *Bathyarchaeota*.

**Figure 3.**
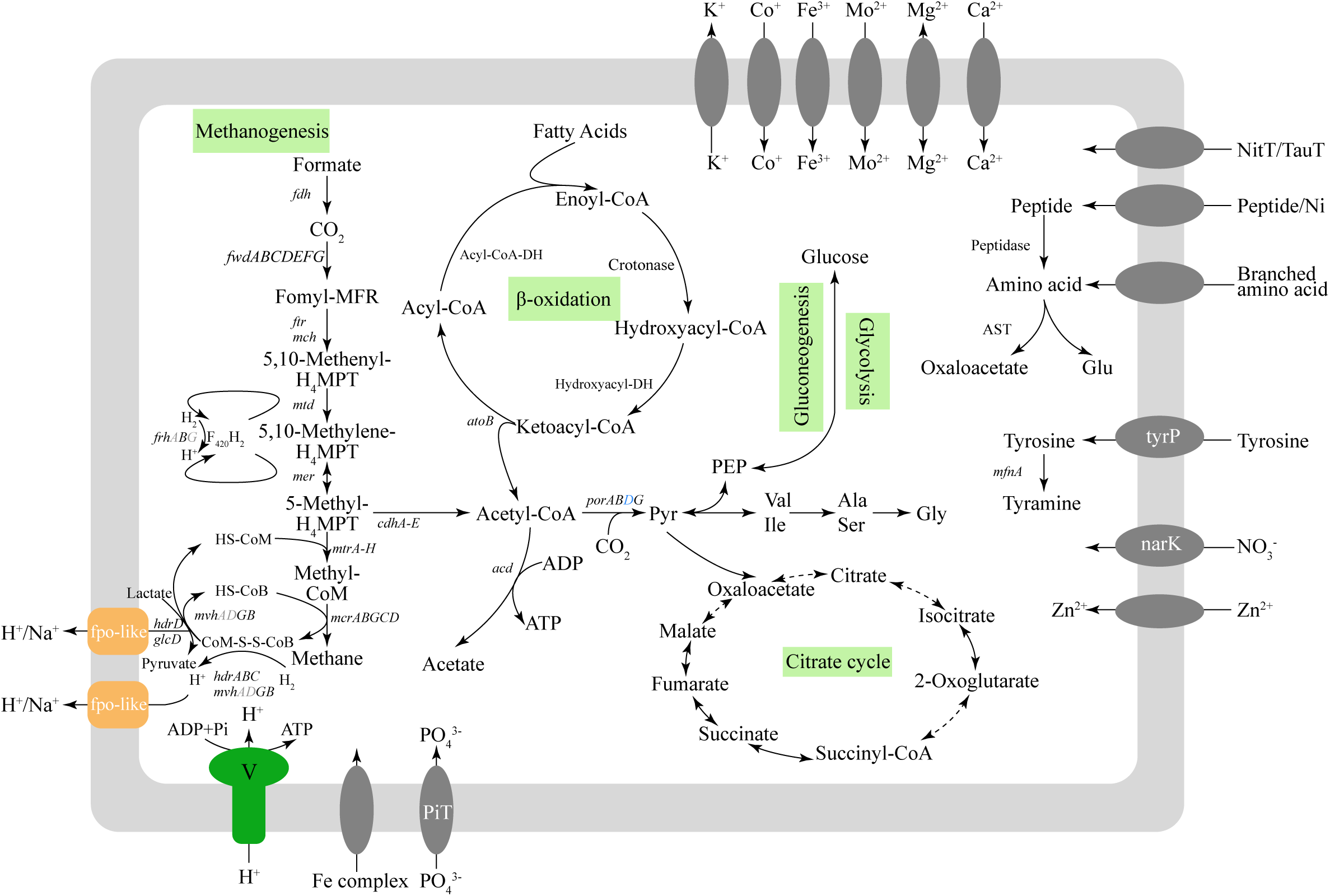
Reconstructed metabolism of the Cren_UShs and Cren_JZhs. The metabolic potentials of crenarchaeotal methanogens are primarily inferred based on the annotation results of IMG/ER. Genes found in both Cren_UShs and Cren_JZhs (black), only found in Cren_UShs (blue) are indicated, or missing from both genomes (gray). Dashed black arrows indicate genes that are missing or unknown in Cren_UShs and Cren_JZhs.

Genes involved in the reduction of heterodisulfide (CoM-SS-CoB) formed during the final step of methanogenesis are identified in these two MAGs. The heterodisulfide reductase (Hdr) and F_420_-non-reducing hydrogenase (Mvh) for the conventional CoM-SS-CoB are found in Cren_UShs and Cren_JZhs, while genes of *mvhA* and *mvG* were not identified in these two archaea genomes. Besides that, the genes for HdrD are in three copies in both Cren_UShs and Cren_JZhs, and two copies in each MAG are co-located with genes with flavin adenine dinucleotide-containing dehydrogenase *(glcD)*, which is in line with the previous reported methane-metabolizing archaea within *Verstraetearchaeota* and *Bathyarchaeota*. The *glcD* coupled *hdrD*, therefore, may be adopted in these newly recovered archaea to reduce CoM-SS-CoB with carboxylate utilization (Evans *et al*., 2015; Hocking *et al*., 2014; Vanwonterghem *et al*., 2016). In addition, homologues of the membrane-bound NADH-quinone oxidoreductase (Nuo), which might be the F420-methanophenazine oxidoreductase (Fpo), are identified in Cren_UShs and Cren_JZhs and perform the re-oxidizing reduced ferredoxin. However, several subunits of fpo are not identified in these two MAGs (Supplementary Table S3) which might be complemented by energy-conserving hydrogenase B (Ehb) (Thauer *et al*., 2008).

## Other metabolic traits and environmental distribution

In addition to the methanogenic metabolism, genes for peptide, amino acid transporter and peptidase were found in Cren_UShs and Cren_JZhs, suggesting these organisms have potential for the degradation of extra-cellular peptide/amino acid (Fig. 3). It is similar to the members in the phylum *Verstraetearchaeota* that Cren_UShs and Cren_JZhs appears to be capable of using sugar to generate acetyl-CoA via the Embden-Meyerhof-Parnas (EMP) (Vanwonterghem *et al*., 2016). Besides, the presence of genes encoding Acyl-CoA dehydrogenase, 4-hydroxybutyryl-CoA dehydratase and 3-hydroxyacyl-CoA dehydrogenase (Supplementary Table S4) suggests that β-oxidation process may be adopted in these newly recovered methanogens (Laso-Perez *et al*., 2016). However, the genes for H_2_-dependent methylotrophic methanogenesis that found in both *Bathyarchaeota* and *Verstraetearchaeota* populations are missing from Cren_UShs and Cren_JZhs genomes, indicating these *Crenarchaeota-like* methanogens conserve energy using the hydrogenotrophic methanogenesis. Comparing with the methanogens of the phyla *Verstraetearchaeota* and *Bathyarchaeota*, this study is the first report of hyperthermophilic methanogens that were outside the phylum Euryarchaeota, and genes encoding DNA reverse gyrase may afford thermal protection for these two methanogens even at a low GC content (~44%) (Forterre 2002).

The microbial community of the US hot spring was dominated by crenarchaeotal populations, while the newly recovered methanogenic archaea (Cren_UShs) only accounted for 0.10% and 0.17% of the all recovered population genomes in US80 and US89, respectively (Supplementary Figure S6). The environmental distribution of the newly recovered methanogenic archaea we evaluated by public database searches for *mcrA* genes. The best hits (>80% amino acid identity) to the Cren_UShs *mcrA* gene sequence are mainly found in hot spring samples (Supplementary Table S5). The number of identified *mcrA*-like sequences is as much as 1085 in one dataset of Jinze hot spring, suggesting these novel archaea may play contribute to the methane cycling in some thermophilic systems. One common trait of those samples that contain these methanogens is the circumneutral condition.

In summary, the present study provided genomic evidence that methane metabolism may be conducted in hyperthermophiles within *Crenarchaeota*. The genes for short-chain hydrocarbon oxidation suggest these methane-metabolizing archaea may oxidize short-chain hydrocarbon. However, the ecological roles of Cren_UShs and Cren_JZhs are inferred by the genomic information, which need further physiological and omic studies in the future.

## Materials and methods

### Sampling and sequencing

Water samples were collected from Ulu Slim hot spring in Malaysia (August 4, 2016) at two different points, one was the main spring head (89°C) and another was the secondary spring head (80°C). The US hot spring is a neutral hot spring with the highest temperature of 104°C. For each sample a 1 L water sample of spring water at each spring head was collected and *in-situ* filtered through a 0.45 μm hydrophilic PTFE membrane (Whatman, USA). DNA was extracted from the filtered biomass using Fast DNA Spin Kit for Soil (MP Biomedicals, France). DNA extracts from the 89°C (US89) and 80°C (US80) hot spring water samples were sequenced on the Illumina HiSeq 4000 platform (Illumina, USA) using 150 bp paired-end reads with a 350 bp insert size.

### Sequencing processing and metagenome binning

The US80 and US89 metagenomes were quality filtered and co-assembled using CLC’s *de novo* assembly algorithm (CLC GenomicsWorkbench v6.04, CLCBio, Qiagen) with *k*-mer of 35 and minimum scaffold length of 1 kb, resulting in 47,532 scaffolds with N50 of 5,337 bp, totaling 166.6 Mb. Metagenome assembled genomes (MAGs) were recovered based on the coverage profile, *k*-mer signatures and GC content (Albertsen *et al*., 2013). The completeness of recovered MAGs was assessed using CheckM (Parks *et al*., 2015). All retrieved MAGs were firstly uploaded to Kyoto Encyclopedia of Genes and Genomes (KEGG) GhostKOALA (Kanehisa *et al*., 2016) for the preliminary reconstruction of metabolic traits, and the MAG contained *mcrA* gene was submitted to the Integrated Microbial Genomes Expert Review (IMG-ER) (Markowitz *et al*., 2009) system for genome annotation. The taxonomy of the putative methanogenic MAG was evaluated by GenomeTreeTK v0.0.3 (https://github.com/dparks1134/GenomeTreeTK) using classify_wf workflow. The relative abundance of recovered MAGs was estimated based on the percentage of mapped reads of a given MAG to all the recovered MAGs, which was normalized by the genome size of bins.

### Data mining of newly recovered *mcrA* sequences

A total of 332 datasets were downloaded from public database that included metagenomes from anaerobic digesters (78), hot springs (81) and hydrothermal vents (222). Ublast (Edgar 2010) was used to screen these metagenome datasets (*e*-value of 1e-5 and accel of 0.5) against an *mcrA* sub-database which contains the newly recovered and other reported *mcrA* sequences. The reads with a bitscore >50 and a top hit to one of the newly recovered *mcrA* sequences were collected, and further filtered with similarity to retain the hits with ≥80% similarity. Datasets that contains novel *mcrA* gene sequences were subsequently assembled to recover new MAGs and *mcrA* genes.

### Genome and functional gene trees construction

A concatenated set of 56 archaeal ribosomal proteins (RPs) were used to construct the genome tree. RPs were extracted from 704 publicly available archaeal genomes and the two newly recovered *Crenarchaeota*-like MAGs using hidden Markov models (HMMs). These amino acid sequences were aligned using MUSCLE (Edgar 2004), and a genome tree constructed from the concatenated alignments using FastTree (v2.1.7) under the WAG and GAMMA models (Vanwonterghem *et al*., 2016). The tree was visualized and decorated using iTOL(Letunic and Bork 2016), and rooted using the DPANN superphylum as an outgroup.

The key genes for the enzymes involved in methanogenic metabolism, i.e., McrA, McrB and McrG, were extracted from the newly recovered MAGs and searched against the NCBI nr database (March, 2018) using Blastp (Camacho *et al*., 2009) to identify similar homologues. Complete amino acid sequences from the phyla *Bathyarchaeota*, *Methanomassiliicoccales, Methanobacteriales, Methanococcales, Methanosarcinales, Methanocellales, Methanomicrobialesand* and *Verstraetearchaeota* were used to generate a phylogenetic tree construction using FastTree (v2.1.7). The functional gene trees were rooted using the *Bathyarchaeota*.

### Functional genes and genome comparison

Open reading frames (ORFs) for the contigs/scaffolds that contained *mcr* genes were predicted into using Prodigal (Hyatt *et al*., 2010) and compared using Blastp. The genoPlotR (Guy *et al*., 2010) R package of was used to visualize the gene loci co-located with the *mcr* genes.

Metabolic comparison across archaeal genomes and the newly recovered MAGs were performed based on the classified KO groups. Briefly, annotation results of all archaeal genomes in the KEGG database (Release 80.1) were firstly collected and combined with the four high-quality *Verstraetearchaeota* genomes (V1-V4) and the newly recovered *Crenarchaeota-like* MAGs, resulting in 167 archaeal genomes. The KO table was then summarized based on the KEGG (161 archaeal genomes) and IMG/ER annotation results *(Verstraetearchaeota* genomes and newly recovered MAG) and further used in a principle component analysis (PCA).

Tetranucleotide frequency-principal components analysis (TNF-PCA) across the newly recovered MAGs and *Verstraetearchaeota* population genomes was used to identify the taxa distance, a genome within the order *Thermoproteales (Thermofilum uzonense)* was selected as outgroup genome. The scaffolds/chromosome of selected genomes were shredded into sequences with length of 5 kb (4 kb overlapping). Nucleotide frequency of each sequence (>2 kb) was calculated with oligonucleotide word size of four. The summarized 4-kmer table of compared genomes was used in PCA analysis. The average amino acid identity was calculated by CompareM (https://github.com/dparks1134/CompareM).

## Acknowledgements

This work was supported by Hong Kong General Research Fund (172057/15E). Mr. Yulin Wang, Mr. Lei Liu wish to thank the University of Hong Kong for the postgraduate scholarships. Dr. Kian Mau Goh appreciated Universiti Teknologi Malaysia Research Grants (Grant No. 16H89, & 14H67).

## Data availability

The sequences have been deposited at the NCBI Sequence Read Archive under BioProject no. PRJNA449277.

